# A neural decoder for learned vocal behavior

**DOI:** 10.1101/193987

**Authors:** Ezequiel M. Arneodo, Shukai Chen, Vikash Gilja, Timothy Q. Gentner

## Abstract

Brain Machine Interfaces (BMIs) hold promise to restore impaired motor function and, because they decode neural signals to infer behavior, can serve as powerful tools to understand the neural mechanisms of motor control. Yet complex behaviors, such as vocal communication, exceed state-of-the-art decoding technologies which are currently restricted to comparatively simple motor actions. Here we present a BMI for birdsong, that decodes a complex, learned vocal behavior directly from neural activity.

---

State of the art BMIs succeed at decoding behavioral intention from brain activity by mapping features of neuronal ensemble activity onto a motor space ^1,2^. Yet the these motor spaces are confined by current technologies to rather simple actions. To prototype a decoder of complex, natural communication signals from neural activity, we capitalize on two aspects of birdsong, a powerful animal model for vocal learning that shares many features with humans speech ^3,4^. First, birdsong is temporally structured (like human speech); this temporal patterning can be built into a decoder using a recurrent neural network ^5^. Second, the biomechanics of birdsong production are well understood; this enables us to employ a biophysical model of the vocal organ that captures most of the complexity of the song and reduces it to a lower dimensional parameter space ^6,7^. By combining these techniques, we decode realistic synthetic birdsong directly from neural activity.

Our decoder interfaces with the sensory-motor nucleus HVC (used as proper name), where neurons generate high-level motor commands that shape the production of learned song. Adult zebra finches (Taeniopygia guttata) sing renditions of a stereotyped motif (a sequence of 3-10 syllables), whose temporal and/or motor structure is thought to be encoded in the activities of two major types of HVC neurons (Fig. 1a) ^8–13^. We implanted 16/32 site Si-probes in male, adult zebra finches and recorded simultaneously their song and neural activity in HVC; then we used these data to train a long-short-term memory network (LSTM ^5^) to translate neural activity directly onto song. The goal of the network is to predict the spectral components of the song at a time bin t_i_, given the values of neural activity features over • previous time bins 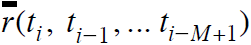 (Fig. 1d-f). The neural activity is fed as a matrix comprising mean firing rates in each time bin, of each putative single/multi-unit automatically sorted from the recordings ^14^ (32/64 clusters); the spectral components of the song are represented by the power across 64 log-spaced frequency bands. For each session (day), we separate the 70-110 renditions of a motif the bird sang. We then train the LSTM network to find the neural-to-song spectrum mappings, and decode the corresponding spectral components from a test set of neural activity to finally recover waveforms of synthetically generated song motifs. We employed several methods to avoid overfitting. First, the order in which each pair of neural feature window/target was presented to the network was randomized, so that the predictions of the spectral components at each time point are independent; second, we used standard techniques such as L2 weight regularization, dropout and early stopping (see Methods). We also employed two different procedures to produce the training/validation and test sets. For motif-wise training/decoding, we split the data into non-overlapping sets of song motifs (reserving 10% for testing), and trained/decoded on these sets. Alternatively, for piece-wise training/decoding we trained the network leaving a fraction of each motif out of the training set, so that the network was tested on an entirely novel song segment); we repeated this procedure using non-overlapping test segments to obtain complete decoded motifs (see Methods).

**Figure 1.**
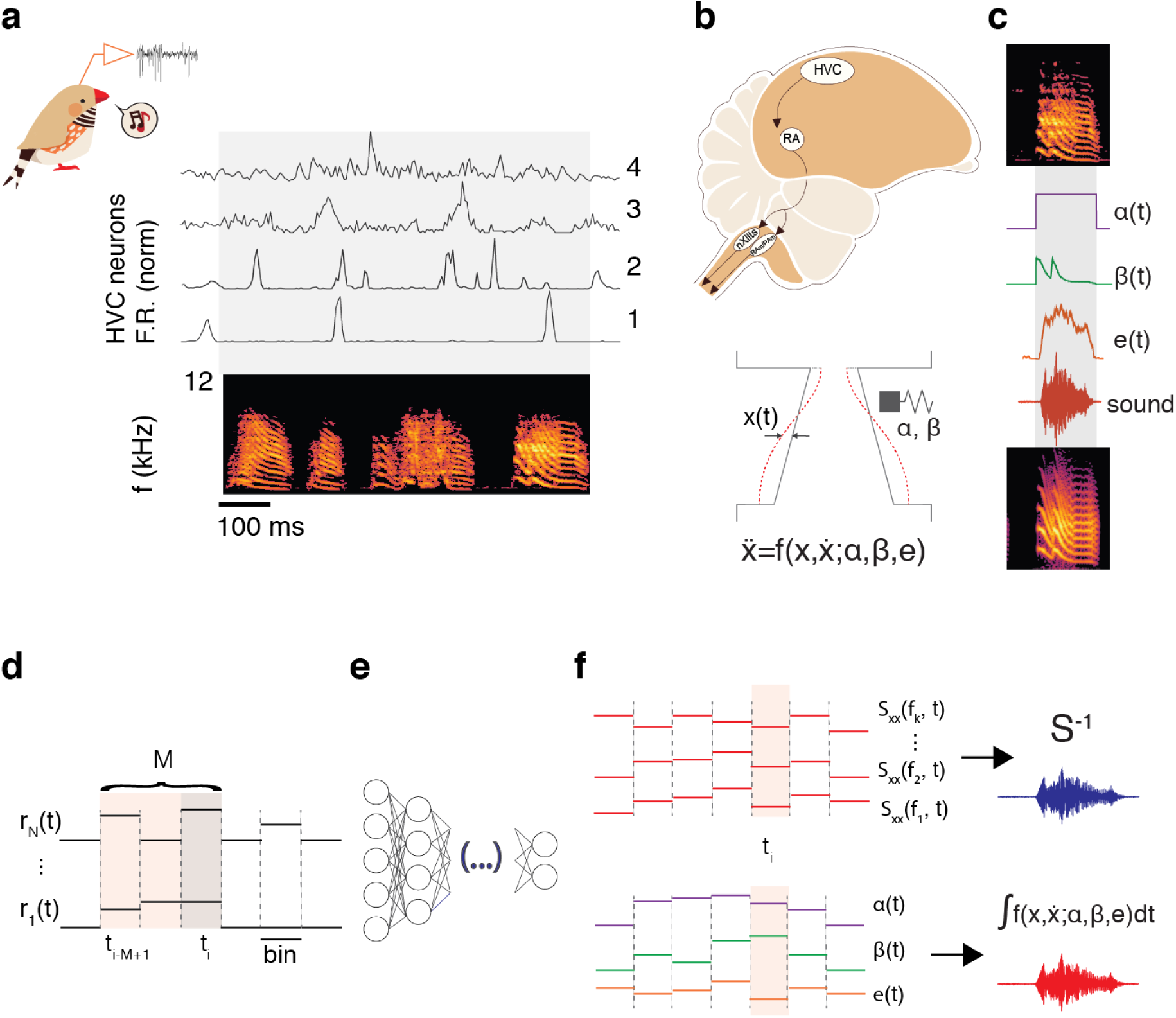
A neural network based decoder to synthesize birdsong from premotor neural activity. **(a)** Neural activity is collected from awake-singing animals. Sorted, extracellularly recorded single/multi-units show different degrees of singing-related sparseness, robustness and spiking precision (4 example clusters, top traces: normalized mean firing rate over 70 aligned to renditions of the bird’s motif, spectrogram). **(b)** Downstream from HVC, the posterior motor pathway nuclei (nXII, RAm/PAm) control the muscles driving sound production ^20^. Syringeal and respiratory muscles act coordinately to modulate the flow of air through sets of labia and produce sound ^15^. The complex labial motion is captured by the equations of a nonlinear oscillator ^17^; parameters that define acoustic properties of the sounds are surrogates of the activities of syringeal and respiratory muscles ^6^. **(c)** To reproduce a particular vocalization (top) from the biophysical model, we fit the parameters (middle {α(t), β(t), e(t)}) such that upon integration, the synthetic song (bottom) matches the pitch and spectral richness (see Methods). **(d)** The input of the decoder neural network is an array with the values of a set of neural features (spike counts of sorted units/multi-units) over a window of M previous time steps. **(e)** The hidden layers of the decoder network are composed of densely connected LSTM cells. **(f)** When training/decoding directly the spectral features of the song, the output of the network is a vector of powers across a range of frequency bands at a given time; the decoded spectral slices are then inverted to produce synthetic song (top). When training/decoding through the biophysical model, the output of the network at a given time is a 3-dimensional vector of parameters (as illustrated in c); the equations of the model are then integrated with these values to produce synthetic song (bottom).

The synthetic neurally decoded songs sound similar to the intended bird’s own song (BOS) (Supplementary files A1, A2; Fig 2a, b; other birds in Fig. S1). We quantitated this similarity by computing the spectrogram root-mean-square-error (sRMSE) between each decoded motif and its corresponding BOS. For reference, we compute the sRMSE between all pairs of motifs sung by the bird in the session. As a control we show the sRMSE between each motif (BOS and synthetic) and a set of motifs from 47 conspecific birds (CON). By this measure, the decoded songs are remarkably like the highly stereotyped bird’s own song: the sRMSE of the decoded songs are within the range of intrinsic BOS variability, and mirror the dissimilarity between BOS and CON (Fig 2d).

**Figure 2.**
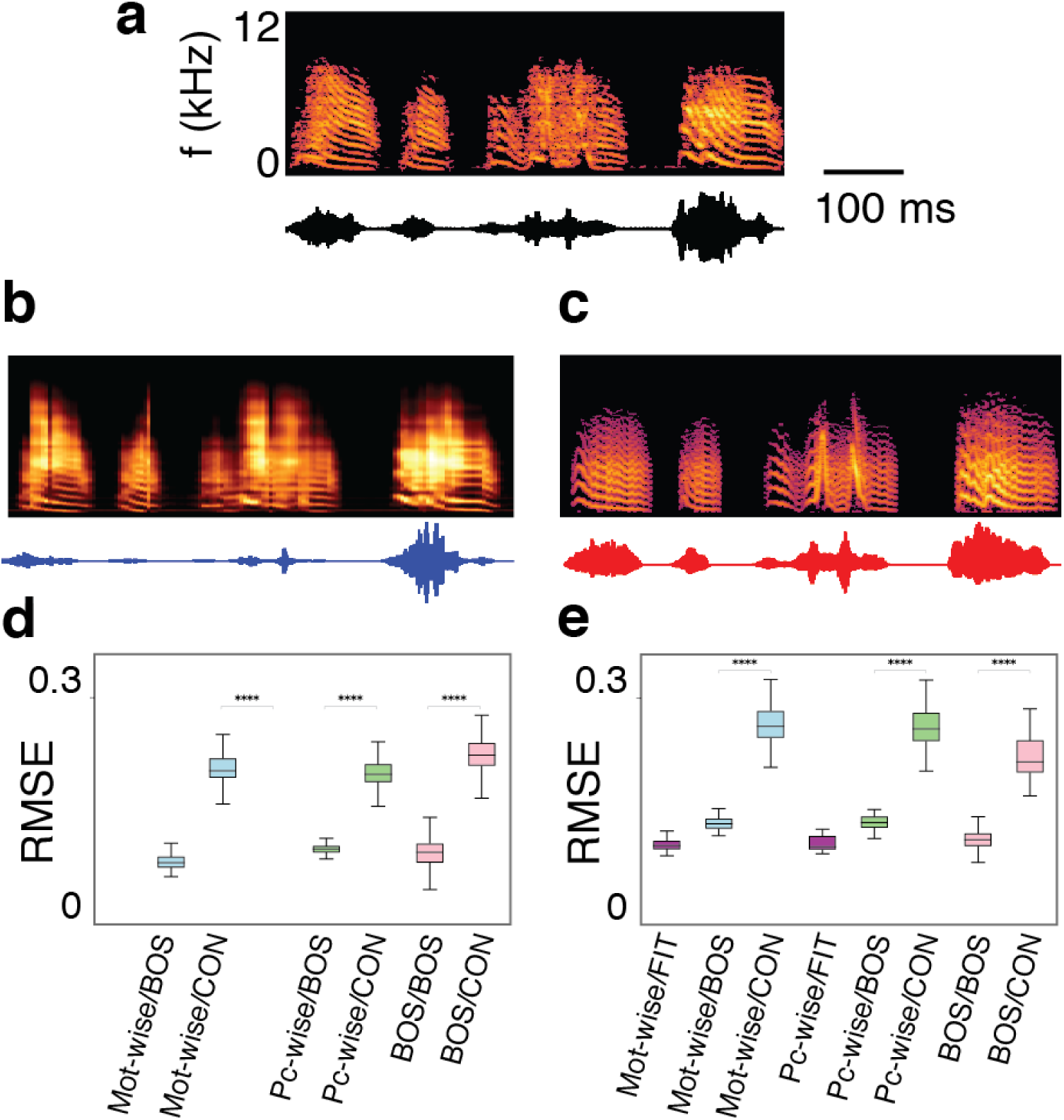
Song decoded from premotor neural activity is similar to the recorded bird’s own song. **(a)** Spectrogram and waveform of one rendition of a bird’s motif (z007). **(b)** Motif generated by decoding spectral components from neural activity and inverting the spectrogram. **(c)** Motif generated by decoding biophysical model parameters from neural activity and integrating the model. **(d)** Performance of the LSTM decoder trained on spectral components (session z007-09-09). Each boxplot summarizes all pairwise spectrogram-RMSE comparisons of: each decoded motif and its corresponding target when training the network with entire motifs (Mot-wise/BOS), each decoded motif and its target when training the network with pieces of motif (Pc-wise/BOS) and a reference provided by different renditions of the bird’s own song (BOS/BOS); also, for control, same measures for renditions of the bird’s motif and motifs of conspecific birds (BOS/CON), decoded motifs and conspecific motifs (Mot-wise/CON and Pc-wise/CON). **(e)** Same as in (d) for the LSTM decoder through the biophysical model (session z007-09-10), with the addition of sRMSE comparisons between each motif decoded from neural activity and its corresponding synthetic motif obtained by integrating the model with the target parameters (Mot-wise/FIT and Pc-wise/FIT)). Sounds represented in panels (a, b, c) correspond to supplementary audio files (A1, A2, A3). (p<1e-10; two-sided Mann-Whitney test).

We can simplify the decoding problem by factoring in the biomechanical apparatus that stands between the motor commands and the vocal output. The syrinx comprises two sets of labial folds that oscillate and modulate the airflow to produce sound ^15^ (Fig. 1b). The dynamics of these labia can be modeled after the motion equations of a nonlinear oscillator, in which the features of the sounds produced are determined by only two parameters ^6,16,17^, which represent the physiological motor instructions driving the syrinx (the respiratory and syringeal muscular activities) ^6^. This model can produce synthetic vocalizations in hard real time ^7^, and vocalizations generated by it are realistic, in the sense that their playback to the asleep/anesthetized animal elicits auditory responses in HVC neurons that match the high selectivity observed in response to BOS ^9^. This model allows us to reduce the dimensionality of the decoder’s target space: for each vocalization recorded, we find the parameters that produce, upon integration of the differential equations of the model, the closest match in pitch, spectral richness and amplitude (Fig. 1c, S5, S6). Thus we represent each song segment as a time series in a 3d parameter space. We then train the network to generate the parameter values that correspond to a set of spiking activities (following the same mechanics as before) (Fig. 1d-f), and finally feed these parameters to the biophysical model and integrate it to produce sound.

This biophysical model-based decoder yields vocalizations that sound similar to the natural ones (Supplementary files A1, A3). The sRMSE between BOS/decoded songs is significantly smaller than the control (BOS/CON), even though the network does not target similarity in the spectral components but in the parameters driving the model (Fig. 2e; Fig. S6 for performance on decoding the parameters).

The decoders we present here can lead to applications in real-time BMIs. The computations involved in decoding parameters (spectral/biophysical) and synthesizing song thereafter are prone to real time (inverting spectrograms/integrating the biophysical model). We can also enhance the efficiency of the neural feature representation by skipping the computational burden of spike sorting: instead, we train/test the decoders using suprathreshold sharp events (unsorted spikes) (Fig. S4, S5). We are also able to decode the spectral features up to 30 ms into the future, thus allocating time for computations between the readout of the activity and the synthesis of the intended song (Fig. S3). Moreover, the simplification achieved by the biophysical model allows us to implement the decoder using a lightweight feed-forward neural network (FFNN) (Fig. S6).

We have demonstrated a BMI for a complex communication signal, using an animal model for human speech and dopaminergic motor learning ^4,18^. Our decoder unlocks access to new models and experiments directed at understanding how neuronal activity is transformed into natural actions. Furthermore, it provides a biomechanic nexus between intention and complex motor output, enabling the question of how the peripheral effectors of motor behavior shape the neural coding of motor intention ^19^. Because the BMI interfaces with a premotor area that is analog to the human vocal motor cortex ^4^ and the computation blocks involved are implementable in real-time, our approach also provides a valuable proving ground for biomedical speech-prosthetic devices.

## Methods

### Subjects

All procedures were approved by the Institutional Animal Care and Use Committee of the University of California (protocol number S15027). Electrophysiology data was collected from n=3 adult (>120 dph) male zebra finches. Birds were individually housed for the entire duration of the experiment and kept on a 14-h light-dark cycle. The birds were not used in any other experiments.

### Neural and audio recordings

We used 4-shank, 16/32 site Si-Probes (Neuronexus), PEDOT-coated in house. We mounted the probes on an in-house designed, printable microdrive and implanted them targeting nucleus HVC. Audio was registered with a microphone (Earthworks M30) connected to a preamplifier (ART Tube MP). Extracellular voltages and pre-amplified audio were amplified and digitized at 30KHz using an intan RHD2000 acquisition system, open ephys and custom software.

### Song detection

A template matching filter written in python was used to find putative instances of the motif, and then curated manually to rule out false positives.

### Spike sorting

Spikes were detected and sorted using Kilosort; details of the procedure can be found in ^14^. The number of clusters was set to 32 or 64 (twice the number of channels of the probe), and we did not perform a post-hoc instance of manual curation, splitting or merging after the initial automatic splitting.

### Supra-threshold event detection

We wrote scripts in python to detect spiking events in each channel. First, the RMS of each channel was estimated using a running window, over a period of time that ranged from minutes to an hour. Then, events that deviated in absolute value more than a number of RMS (2.5-5.5) were detected using the package peakutils (min_distance=0.5ms).

### Dataset preparation

#### Neural activity features

With all 64 clusters spike-sorted, we extracted spike counts within each motif and collapsed them into 4.3ms (128 samples at 30,000 samples/second) time bins. The same time bins were also used for target preparation (both spectral features and biophysical features).

#### Spectral features

When training the networks with spectral features, the target at each time step was a vector containing a spectrogram slice (in log power scale). We first generated 1024-band spectrograms (2048 FFT steps) with each motif waveform. Then we mapped these spectrograms onto mel-filtered spectrograms. The mel scale is a perceptual pitch scale that is equidistant judged by listeners ^21^. The conversion between mel scale and frequency we used was first introduced in ^22^:

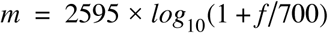

Where *f* is frequency expressed in Hertz. Such scale effectively approximates the auditory system as studies have shown that listeners require an increasingly large interval in normal frequency domain in order to achieve the same pitch difference. In our study, a 64-filter mel scale filter bank was applied to convert the 1024-band spectrograms into 64-band mel-filtered spectrograms, each slice of which was used as target of the model. The mel-filtered spectrograms could be easily reversed back to frequency-based spectrograms and subsequently waveforms in time domain.

### Biophysical model of the vocal organ

#### Model

A model of the zebra finch vocal organ has been previously introduced and explained in detail ^6,7,16^. This model considers mainly a sound source and a vocal tract that further shapes the acoustics of the vocalizations.

The source (syrinx), is comprised of two sets of tissues or labia that can oscillate induced by the sub-syringeal pressure and modulate the airflow to produce sound ^15^. The motion of the labia is represented as a surface wave propagating in the direction of the airflow, that can be described in terms of the lateral displacement of the midpoint of the tissue ^17^. Its mathematical form is the motion equation of a nonlinear oscillator in which two parameters that determine the acoustic features of the solutions are controlled by the bird: the sub-syringeal air sac pressure and the stiffness of the restitution (through the activity of syringeal muscles). In order to integrate the model in real time, a set of equations was found that is computationally less expensive yet capable of displaying topologically equivalent sets of solutions as the parameters are varied ^16^:

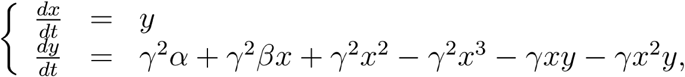

where *x* represents the departure of the midpoint position of the oscillating labia, **γ** is a time scaling factor, and the parameters **α** and **β** are functions of the air sac pressure and the activity of the ventral syringeal muscle, respectively.

The upper vocal tract further shapes the sound produced by the source, determining spectral properties such as the timbre. We used a model for the vocal tract that includes a tube, accounting for the trachea, and a Helmholtz resonator, accounting for the oropharyngeal-esophageal cavity (OEC) ^23^. The pressure at the input of the tract is *Pi*(*t*) = *a* × *x(t) – x(t-τ)*, where *a* × *x(t)* is the contribution to the fluctuations by the modulated airflow, *r* is the reflection coefficient at the opposing end of the tube of length *L* and, with c the sound velocity. We can obtain the pressure fluctuations (the sound) at the output of the system by solving the electrical analog to this acoustic circuit ^6^.

#### Parameter fitting

To fit the parameter series that will lead to reconstruction of the song, we perform a procedure similar to that previously described ^6,9^. Time scale parameter **γ** is set to a value of 23,500; **α** is set to −0.15 during vocalization and 0.15 otherwise, and **β** is set in order to minimize the distance in the (pitch, spectral content) space between the synthesized and the recorded song segments ^6^; the envelope (e(t) in the main text) is obtained by rectifying and smoothing the recorded waveform; the parameters of the OEC were fixed, in the same values as in ^6^. To extract the pitch of the song, we follow a modification of the automatic procedure presented in ^24^, and add a layer of manual curation. When integrating the model, we apply the extracted envelope (e(t)) as an extra multiplicative factor when computing *a* × *x(t)*, since it recovers the amplitude fluctuations that were discarded when reducing the model to its normal form and driving it with the bi-valued parameter **α**.

### Neural network training

Neural network based decoders were coded in python, using Tensorflow and in some cases Keras. They were run on PCs equipped with NVidia GPUs (Tesla k40, Titan Z, and Titan X Pascal).

#### LSTM network architecture

The network has 2 layers of LSTM cells, with 30 units in the first layer and 20 in the second. The output layer has as many linear units as the target space (64 for the spectrogram bands, 3 for the biophysical model parameters). Both LSTM layers utilized 20% dropout and 0.001 L2 regularization during training to prevent overfitting ^25^.

#### Feed-forward Network architecture

The architecture is essentially the same as that of the LSTM network, but it replaces the LSTM layers with three dense layers of relu units ^26^. The first of these layers halves the dimension of the input vector; then each layer halves the dimension of the previous one. All layers utilized 20% dropout and 0.001 L2 regularization during training to prevent overfitting ^25^.

#### Training procedure

We utilized a gradient-based optimizer (Adam/rmsprop ^27^) and mean square error (MSE) as a loss function for LSTM/FFNN. Two training conditions were experimented, referred to as motif-wise and piece-wise training.

*Motif-wise training*: we used 10% of all the motifs for testing and the remaining motifs for training. We made 10 passes using non-overlapping motifs as testing set to have as many decoded examples as number of motifs in the session. In each pass, all of the neural-activity/decoder-target pairs (one per bin) were fed in random order to the network, both when training and decoding.

*Piece-wise* training: we held out a fraction of each motif when training (roughly 3.3%); trained on the complement and generated the song corresponding to the masked fraction; we repeated this procedure tiling the whole motif, and generated entire motifs using segments of data that were novel to the decoder. In both training conditions, 10% of the training set was reserved as validation set for early stopping, where the training session would terminate if validation loss failed to decrease within 5/10 training epochs.

### Song waveform generation

#### Spectrogram inversion

We used the LSEE-STFTM algorithm to invert spectrograms back to audio waves ^28^. The algorithm iteratively estimates a signal from the short-time Fourier transform magnitude (STFTM), through minimizing the mean square error between the short-time Fourier transform (STFT) and the estimated STFT, and subsequently performs STFT on the estimated signal, the magnitude of which will be passed on to the next iteration.

Within each iteration, a signal was approximated using the equation below:

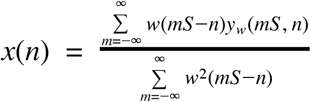

Where *x*(*n*) denotes the estimated signal; *w*(*n*) denotes the analysis window used in STFT. The variable *S* is a positive integer, representing the sampling rate of the STFT. Here, *y*_*w*_ (*mS*, *n*) is the target signal corresponding to *Y* _*w*_ (*mS*, *n*), which denotes the target STFTM, in our case spectrogram powers. To calculate in each iteration, we used a sinusoidal window ^28^

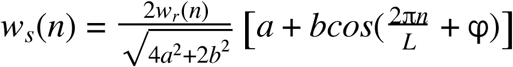

where *L* represents the length of the window. Here, *w*_*r*_ (*n*) is a rectangular window with an amplitude of 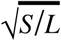 within 0 ≤ *n* < *L* and zero anywhere outside. A modified Hamming window can be obtained by *a* = .54, *b* =–.46, 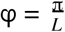 setting After obtaining *x*(*n*) value within each iteration, the STFT of *x*(*n*) was calculated, which was used in place of *Y* _*w*_ (*mS*, *n*) in the next iteration. The squared error between the target STFTM and the estimated STFTM is proven to decrease in each iteration of the algorithm

#### Biophysical model integration

Once the model parameters are predicted by the decoder, they are re-sampled and fed to an ordinary differential equation integrator. Resampling to 30 Khz is performed (with cubic interpolation). A fourth order runge-kutta ODE integrator (custom coded) integrates the equations of the model with a (900 Khz)^-1^time step.

### Performance Evaluation

#### Root Mean Square Error (RMSE)

We evaluated performance of our models using RMSE between each pair of original and predicted spectrogram magnitudes.

#### Spectrogram Normalization

To account for amplitude variations between motifs from different birds, we normalized spectrograms for each bird so that the collection of original spectrograms for each bird had a maximum power of 1 and minimum power of 0:

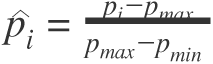

Where *p*_*i*_ is the power of a point on either an original spectrogram or a predicted spectrogram before normalization, while 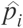 is the normalized power of the corresponding point.*p*_*max*_ denotes the *p*_*i*_ maximum power of the entire set of original spectrograms, while *p*_*min*_ represents the minimum power of the entire set of original spectrograms. With such normalization, we were able to account for variations among motifs from different birds while keeping the variations within motifs from the same bird.

#### Pair-wise performance comparisons

We performed spectrogram-RMSE comparisons within and between different sets of birds (displayd in Fig 2d,e, boxplots for instance). *BOS/BOS:* comparisons provide a baseline of the variability of the bird’s own motifs during the session: spectrogram-RMSE across each pair of renditions of a motif. *Mot-wise/BOS:* spectrogram-RMSE values across each pair of natural motif and the corresponding motif decoded from neural activity, when training/decoding the network *Motif-wise. Pc-wise/BOS:* spectrogram-RMSE values across each pair of natural motif and its corresponding one decoded from neural activity, when training/decoding the network *Piece-wise*.

To provide an additional reference for song variability, we also computed spectrogram-RMSE comparisons in a set of 47 motifs from conspecific birds (other zebra finches; about half of them from our colony and half from other colonies). This produced the sets: *BOS/CON:* spectrogram-RMSE across each BOS rendition and all of the conspecific (CON) motifs. *Mot-wise/CON:* spectrogram-RMSE values across each motif decoded from neural activity and all the CON motifs, when training/decoding the network *Motif-wise. BOS/CON:* spectrogram-RMSE across each BOS rendition and all of the conspecific (CON) motifs. *Pc-wise/CON:* spectrogram-RMSE values across each motif decoded from neural activity and all the CON motifs, when training/decoding the network *Piece-wise*.

## Data availability

The datasets are available from the corresponding authors upon request.

## Materials and code availability

Code, printable hardware and electronic designs developed during this work are available in the following github repositories:

https://github.com/singingfinch/bernardo

<u>https://github.com/zekearneodo/swissknife</u>

## Acknowledgments

We thank K. Perks, B. Theilman, M. Thielk, T. Sainburg, D. Battista for critical discussions and/or input on the manuscript. This work was supported by the U.S. National Institute of Health (Grant R01DC008358), the Kavli Institute for the Brain and Mind (IRG #2016-004), the Office of Naval Research, and a Pew Latin American Fellowship in the Biomedical Sciences (E.A.). The Tesla K40, Titan Z and Titan X Pascal used for this research were donated by the NVIDIA Corporation.

## Author contributions

All of the authors participated in the design of the experiment and the writing of the manuscript. E.A. built the experimental setup and collected the data. E.A. and S.C. analyzed the data. T.G. supervised the study.

## Supplementary Figures

**Figure S1.**
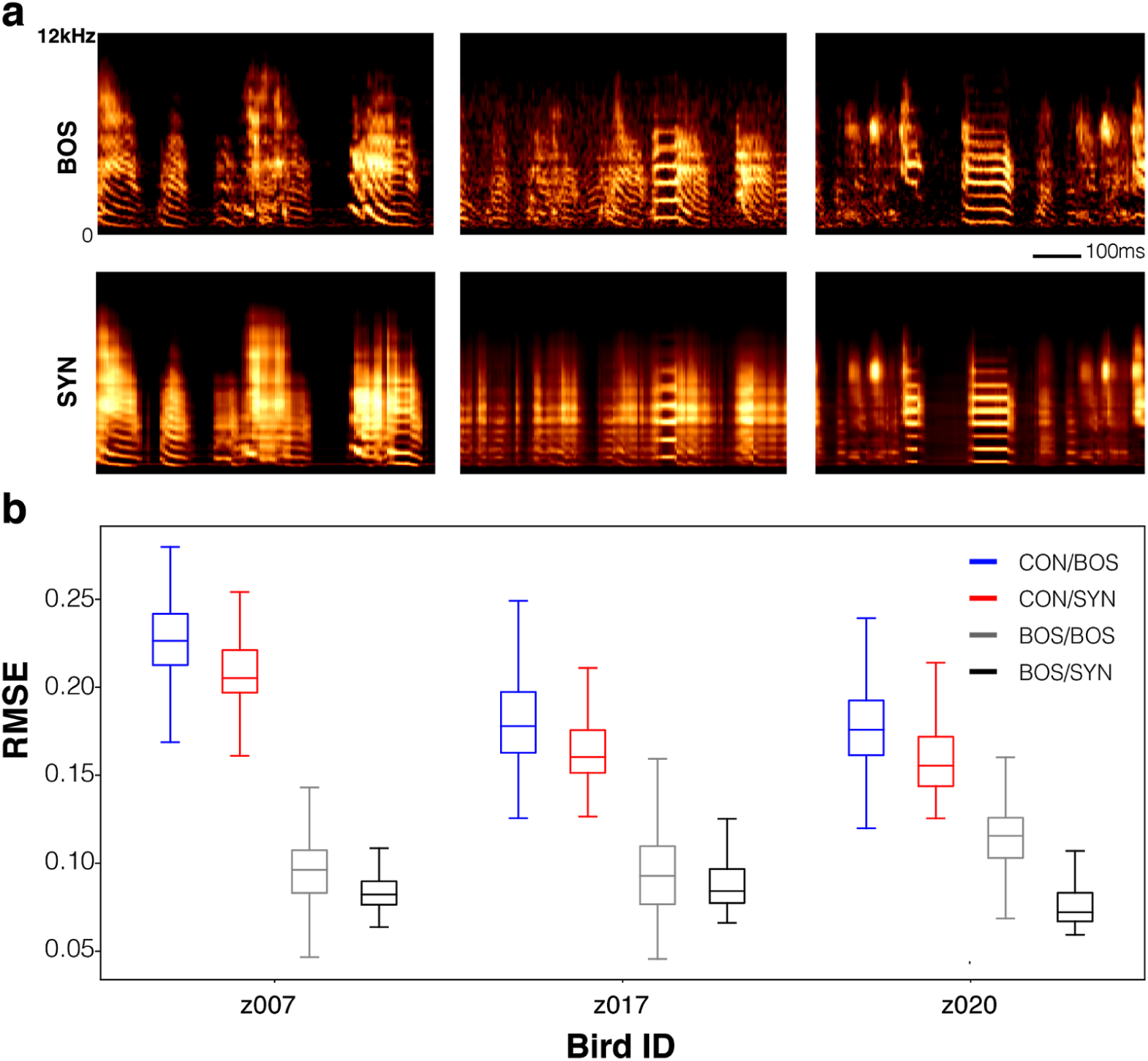
Synthetic motifs from three birds generated by LSTM trained on spectral components. **(a)** Example spectrograms of each bird’s motif (BOS) and the corresponding motif decoded from neural activity (birds z007, z017, z020; one session each). Songs were synthesized by motifwise training/decoding (denoted SYN). **(b)** Performance evaluation, each box plot constructed as described previously for Fig. 1.

**Figure S2.**
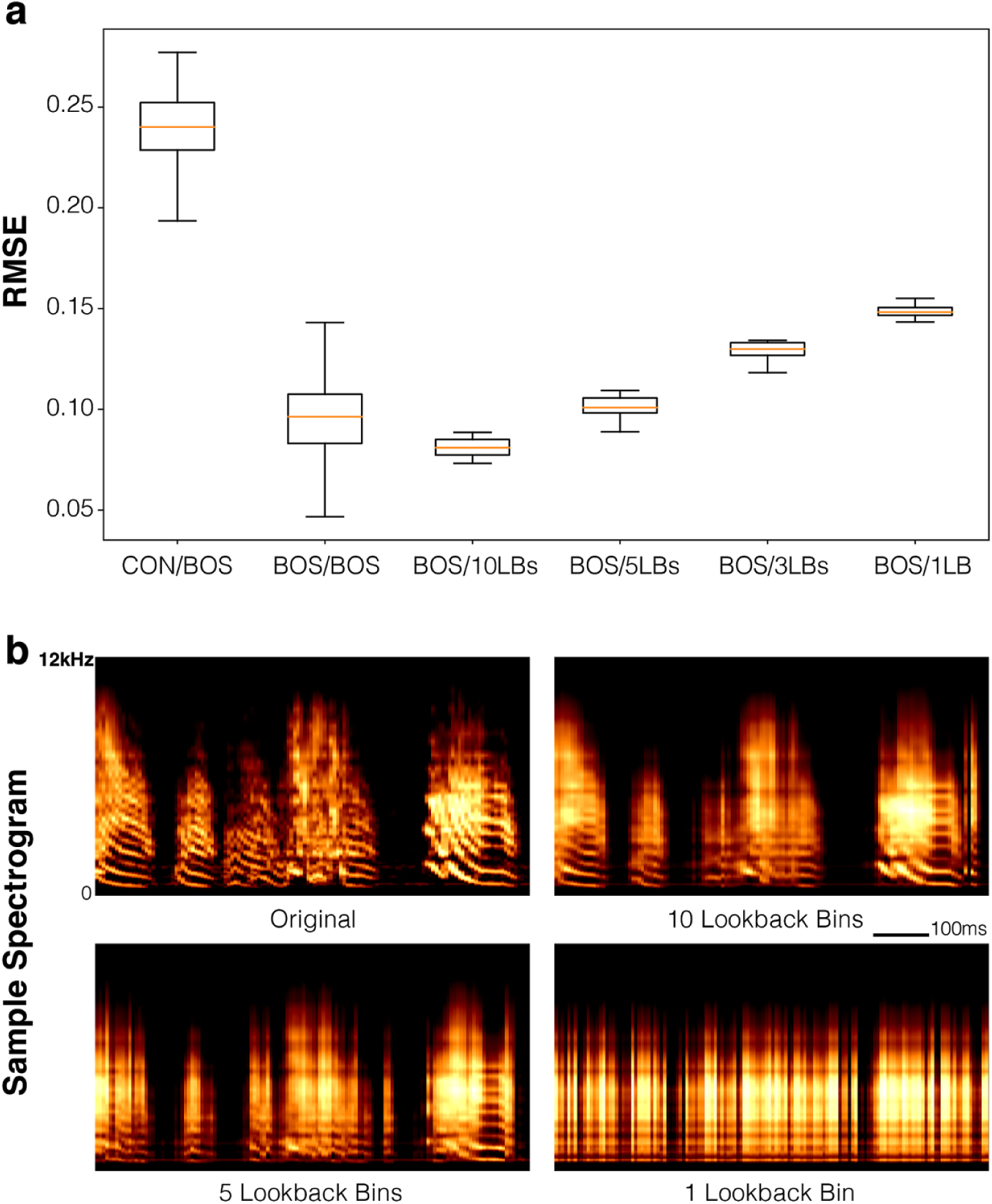
Synthetic motifs decoded from decreasing number of lookback bins (training/decoding through spectral components; motifwise). **(a)** Performance evaluation of synthetic motifs decoded from decresing numbers of lookback bins (LBs) in the window of neural activity fed to the decoder. RMSEs between decoded motifs and their corresponding BOS motifs are compared to RMSEs among the bird’s own songs (BOS/BOS) as well as RMSEs between conspecific motifs and the bird’s own songs (CON/BOS). (b) Example spectrograms of the bird’s own motif and the corresponding synthetic motifs decoded from decreasing numbers of lookback bins.

**Figure S3.**
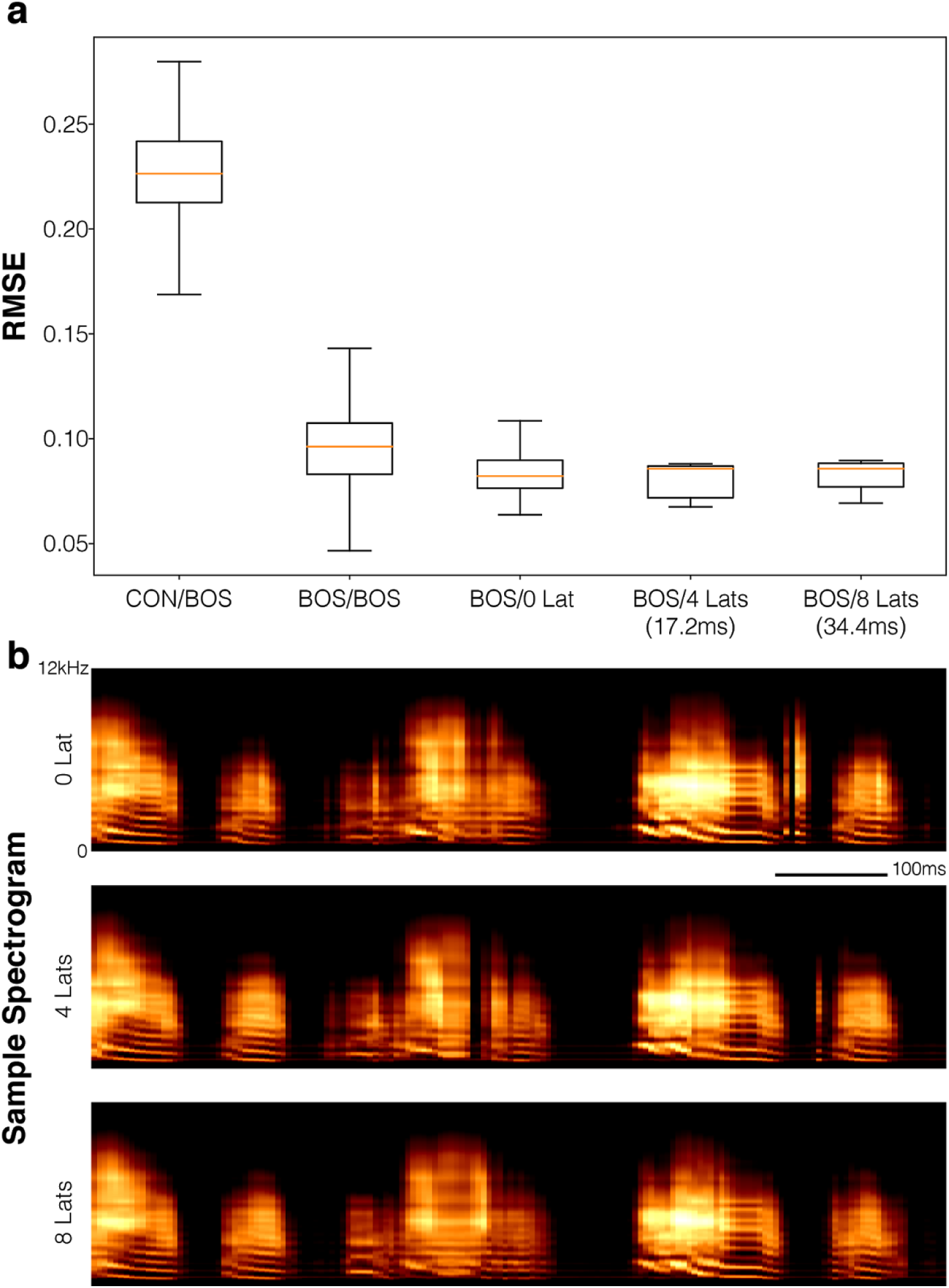
Decoding with a latency between the window of neural features and the target spectral components. **(a)** Performance of motifs decoded with different numbers of latency bins (Lats) (time bins between the right end of the neural featurs window and the target spectral slice). RMSEs between synthetic motifs with different number of latency bins and corresponding BOS motifs, as well as between conspecific motifs and the bird’s own songs (CON/BOS). **(b)** Example spectrograms of synthetic motifs decoded with different number of latency bins (up to 8 bins or 34.4ms of latency).

**Figure S4.**
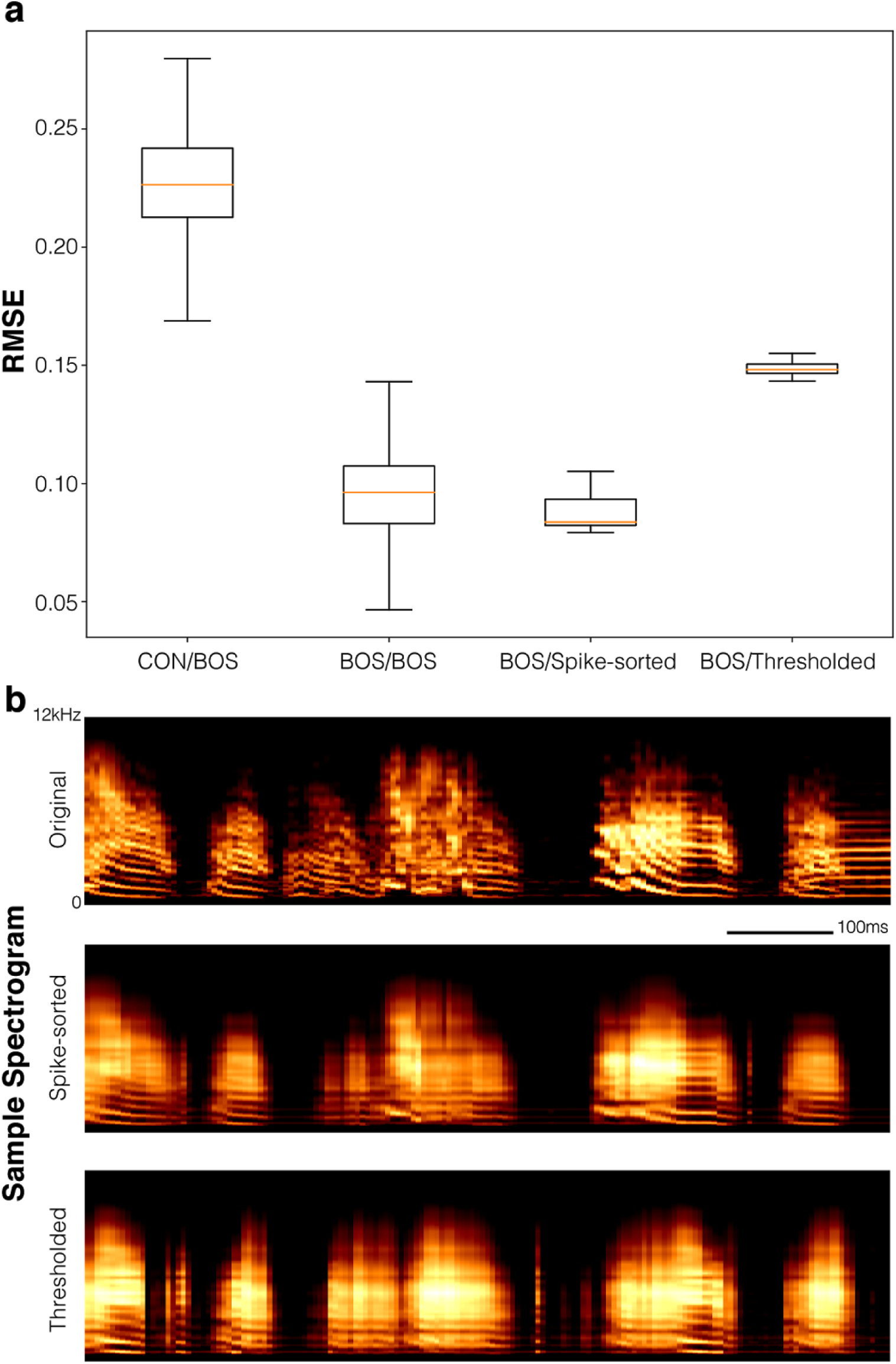
Training the LSTM decoder using spectral features and unsorted spikes (thresholded extracellular activities). **(a)** Performance comparison between synthetic songs decoded from sorted spike counts (BOSSpikesorted) and thresholded activity (BOS/Thresholded). **(b)** Example spectrograms.

**Figure S5.**
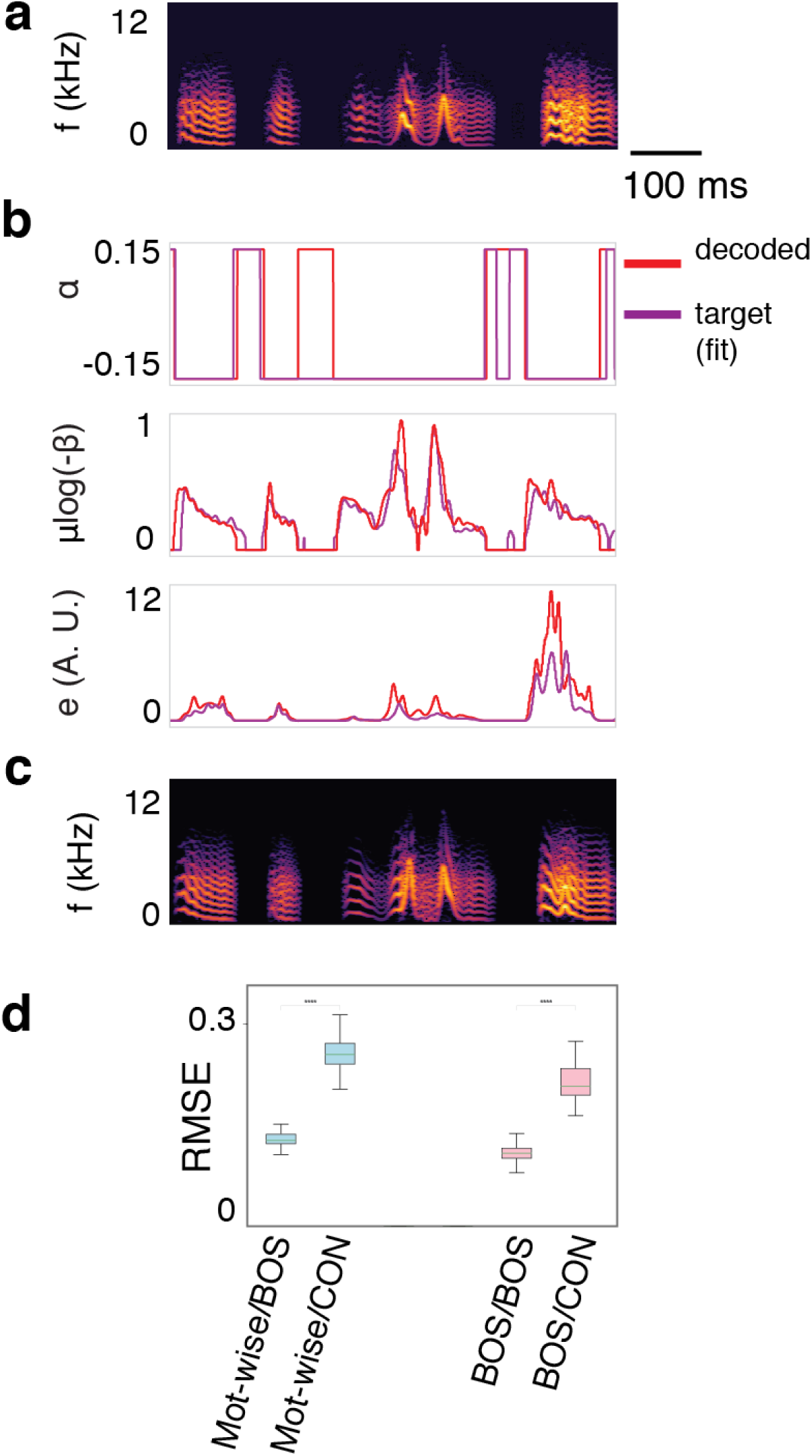
Synthetic song decoded from thresholded extracellular activity (unsorted spikes) and fit detail. **(a)** Decoded motif. **(b)** Decoded parameters of the biophysical model, together with their target (fitted offline to approximate the BOS). **(c)** Synthetic motif corresponding produced by the parameters in (c) (target). **(d)** Evaluation of performance (motif-wise training/decoding). (p<1e-10; two-sided Mann-Whitney test).

**Figure S6.**
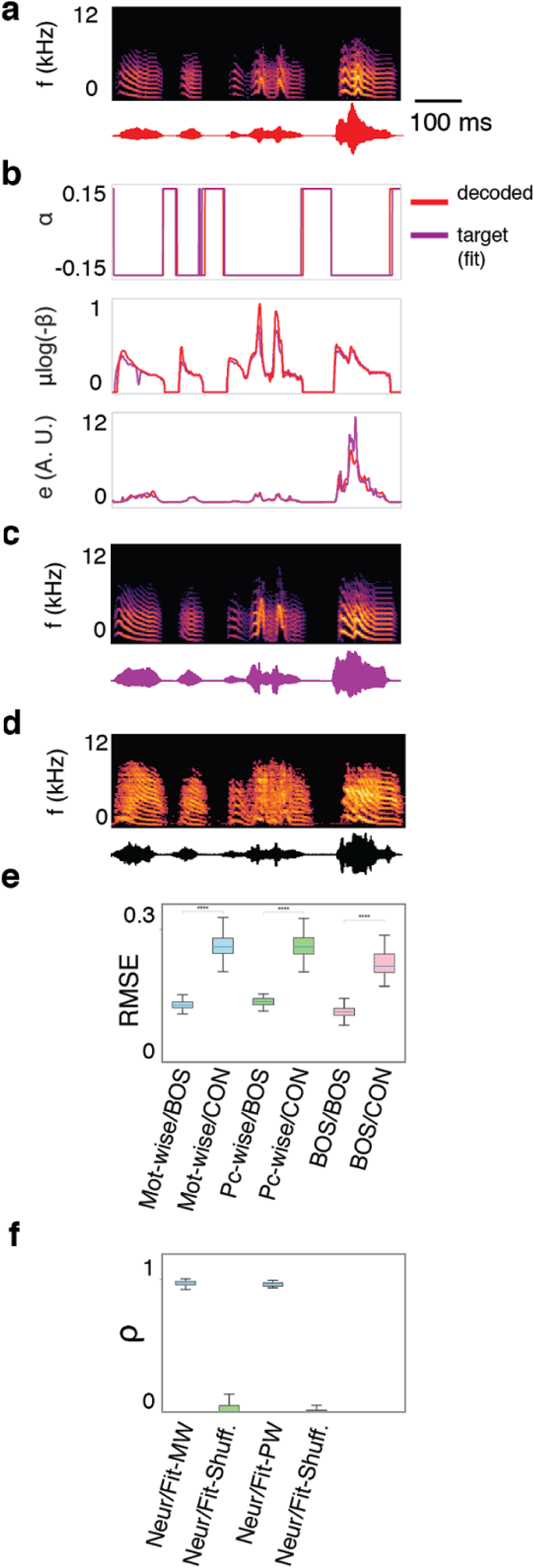
Synthetic song decoded from sorted spikes using a feedforward architecture and fit detail. **(a)** Decoded motif. **(b)** Decoded parameters of the biophysical model, together with their target (fitted offline to approximate the BOS). **(c)** Synthetic motif corresponding produced by the parameters in (c) (target). **(d)** Corresponding BOS. **(e)** Evaluation of performance (same as caption of Fig. 2 in the main text). **(f)** Mean correlation between target and decoded parameters (Neur/Fit) for each motif. Training/decoding condition denoted by MW (motifwise) or PW (piecewise); for reference we also computed the correlation between target and shuffled decoded parameters for each training condition (Neur/FitShuff.). (p<1e-10; two-sided Mann-Whitney test)

**Figure S7.**
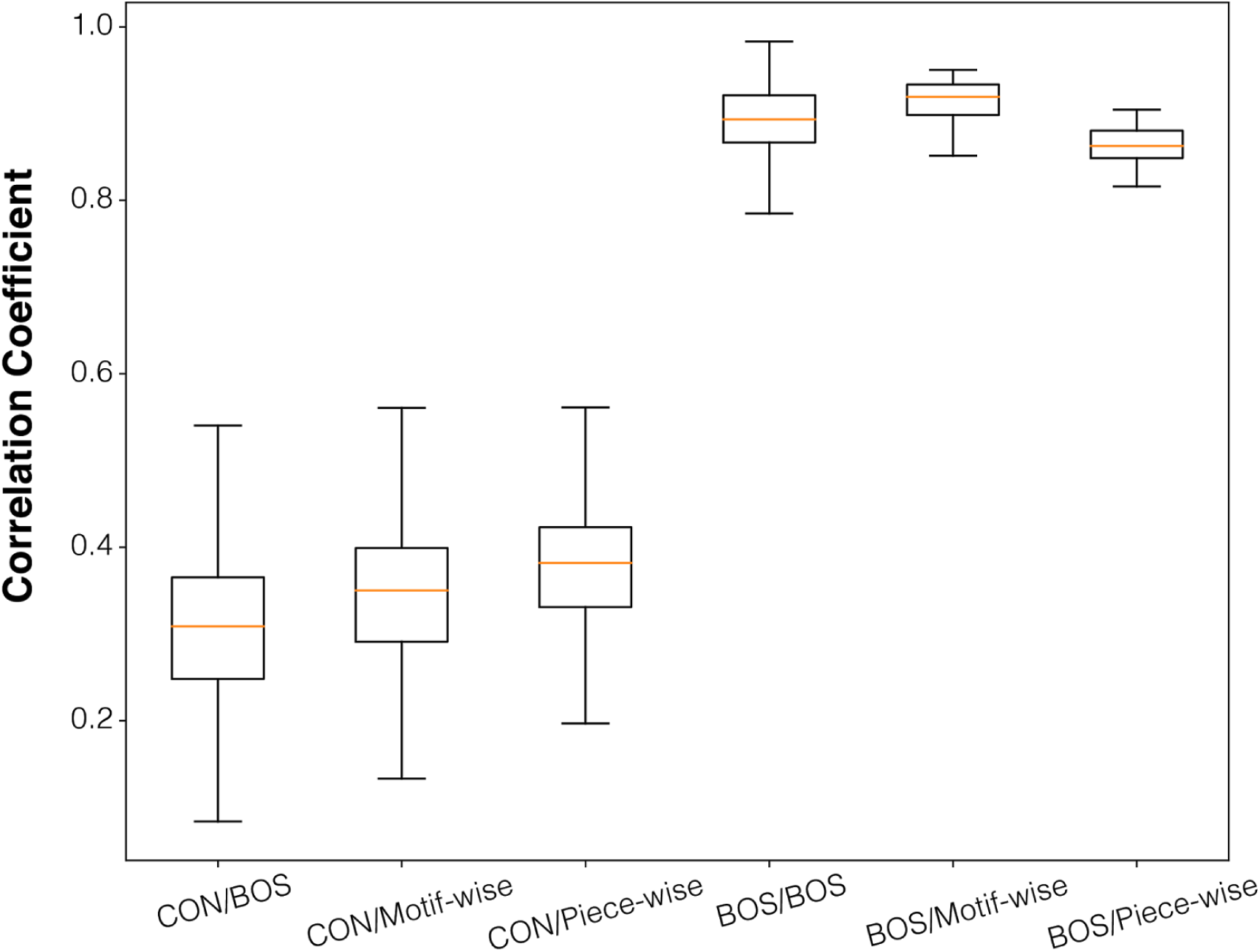
Performance of the LSTM decoder trained on spectral components in terms of correlation coefficient. Correlation coefficients of: conspecific motifs and the bird’s own motif (CON/BOS); conspecific motifs and synthetic motifs decoded by motif-wise training (CON/Motif-wise); conspecific motifs and synthetic motifs decoded by piece-wise training (CON/Piece-wise); all pairs of the bird’s own motif (BOS/BOS); synthetic motifs and the corresponding natural motifs from both motif-wise training (BOS/Motif-wise) and piece-wise training (BOS/Piece-wise).

## Supplementary Files List

**Audio A1:** Example rendition of a motif of bird z007.

**Audio A2:** Example decoded motif of bird z007, training/testing the decoder using spectral features as target (motifwise training).

**Audio A3:** Example decoded motif of bird z007, training/testing the decoder through the biophysical model (motifwise training).

**Audio A4:** Inverted spectrogram of a training example (melspaced spectrogram of the motif), corresponding to audio A3.

**Audio A5:** Example motif of bird z007 synthesized by fitting the parameters of the biophysical model and numerically integrating it (fitted parameters being the actual target of the network when decoding; corresponding to audio file A2).

